# Temporal and spatial population genetic variation in Chilean jack mackerel, *Trachurus murphyi*

**DOI:** 10.1101/2024.08.05.606603

**Authors:** CB Canales-Aguirre, S Ferrada-Fuentes, R Galleguillos

**Affiliations:** Centro i∼mar, Universidad de Los Lagos, Camino a Chinquihue 6 km, Puerto Montt, Chile; Núcleo Milenio INVASAL, Concepción, Chile; Laboratorio de Genética y Acuicultura, Departamento de Oceanografía, Universidad de Concepción, Chile; Doctorado en Sistemática y Biodiversidad, Facultad de Ciencias Naturales y Oceanográficas, Universidad de Concepción, Chile

**Keywords:** genetic homogeneity, jurel, microsatellites, temporal genetic variation

## Abstract

The *Trachurus murphyi* have been studied for population genetic structure for decades, identifying only one large population across the South Pacific Ocean. Although all these studies have extensively examined the spatial genetic pattern, there remains a gap in understanding the potential role of temporality. Our study aims to elucidate temporal and spatial genetic patterns in *T. murphyi* populations in the South Pacific Ocean, examining genetic composition across seasons, including feeding and spawning seasons, where the latter was not previously investigated. Using 10 microsatellite loci, our study confirms a consistent and stable population genetic pattern in *T. murphyi* across its geographic distribution, observed over multiple years and seasons. Furthermore, we identify potential genetic markers for monitoring variability in the species.

## 1. Introduction

The Chilean jack mackerel, *Trachurus murphyi*, is a highly mobile marine fish distributed throughout the South Pacific Ocean (Serra, 1991). It is predominantly found along the coast from Ecuador to Chile in the southeastern Pacific Ocean and near the coasts of New Zealand and the Tasmanian Sea (Bailey, K., 1989; Evseenko, 1987; Serra, 1991). Due to its significant importance for fisheries in this region, numerous studies have investigated population differences within its distribution range, examining various aspects such as life history traits, meristic and morphometric characteristics, microchemistry of otoliths, as well as genetic markers (Canales-Aguirre et al., 2010a; Cárdenas et al., 2009; Cubillos et al., 2008; Evseenko, 1987; Ferrada Fuentes et al., 2023; Galleguillos et al., 2012; Hernández et al., 1998; Horn and Ó Maolagáin, 2021; Oliva, 1999; Serra, 1991; Taylor, 2002; Vásquez et al., 2013).

Numerous studies have been conducted using molecular tools to get insight into the population genetic structure in *T. murphyi* across the South Pacific Ocean. All of them since 80’ with a focus on the integration of population genetic data into a tool for informed decision-making for this species (Canales-Aguirre et al., 2010a; Cárdenas et al., 2009; Ferrada Fuentes et al., 2023; Galleguillos et al., 2012). For instance, Cárdenas et al. (2009) found high genetic diversity using three heterologous microsatellites, and similar results were found by Ferrada Fuentes et al. (2023) using seven microsatellites. These population genetics studies and others have concluded that this species has not shown population genetic structure along its geographic distribution. However, while these studies have extensively examined spatial genetic patterns, there remains a gap in understanding the potential role of temporality. It is important to consider that individuals may migrate to spawning areas before dispersing to feeding areas, where mixing occurs (Glover et al., 2011; Waples, 1998). Therefore, sampling individuals in spawning areas can provide valuable insights and maximize the likely population genetic difference compared with areas where there can mixed.

Our study aims to shed new light on the temporal and spatial genetic composition of *T. murphyi* populations in the South Pacific Ocean. We study genetic patterns over different seasons across a year, including feeding (Winter-Fall) and spawning (Spring-Summer), which were not included in previous studies. Additionally, we aim to identify genetic markers that may serve as indicators of changes in genetic diversity, comparing the expected heterozygosity obtained with previous studies where the same loci were used. Finally, we aim to expand our understanding of population structures in marine species over time and shed light on the evolutionary processes within *T. murphyi*.

## 2. Materials and Methods

### 2.1. Sample collection areas

We collected a total of 852 individuals of *Trachurus murphyi* in 2011-2012 from 18 locations across the coastal areas of the South Pacific Ocean (i.e., Perú, Chile and New Zealand; Fig. 1) during fall-winter (_FW, n= 538) and spring-summer (_SS, n= 314) seasons. The fall-winter season (March to August) corresponds to the main period where fishery vessels catch Chilean jack mackerel; hence, there are no biological reproductive closures. Also, this period corresponds to the feeding time so individuals disperse from their main reproductive areas to several points along the coast and ocean zones. The fall-winter season included 12 locations: six from Perú, five from Chile, and one from New Zealand. The fall-winter locations were aggregated as follows: PePN_FW (Pimentel), PeTCH_FW (Chimbote and Ancon), PePa_FW (Cañete, Tambo de Moras and Olleros) in Perú; ChNo_FW (Iquique and Mejillones), ChCo_FW (Coquimbo), ChThco_FW (Talcahuano coast), ChChoc_FW (Chiloé oceanic) in Chile; and NZ_FW in New Zealand. The spring-summer season (October to February) corresponds to the spawning season, where fishery vessels are forbidden to catch Chilean jack mackerel (Serra, 1991) because of biological reproductive closures. This season, individuals move from feeding to their main reproductive areas (Serra, 1991). The spring-summer season included six locations: one from Perú, four from Chile, and one from New Zealand. The spring-summer locations were aggregated as follows: PeTCH_SS (Chiclayo) in Perú; ChNo_SS (Iquique), ChCo_SS (Coquimbo), ChThco_SS (Talcahuano coast), ChChoc_SS (Puerto Montt) in Chile; and NZ_SS in New Zealand. All previous genetic studies conducted on *T. murphyi* have collected samples from the fall-winter season because accessibility is easier. Here, we included the spawning season of Chilean jack mackerel samples to enhance the robustness in delineating population genetic structure (Glover et al., 2011), and the feeding season, anticipating the mixing of individuals from different areas.

**Figure 1.**
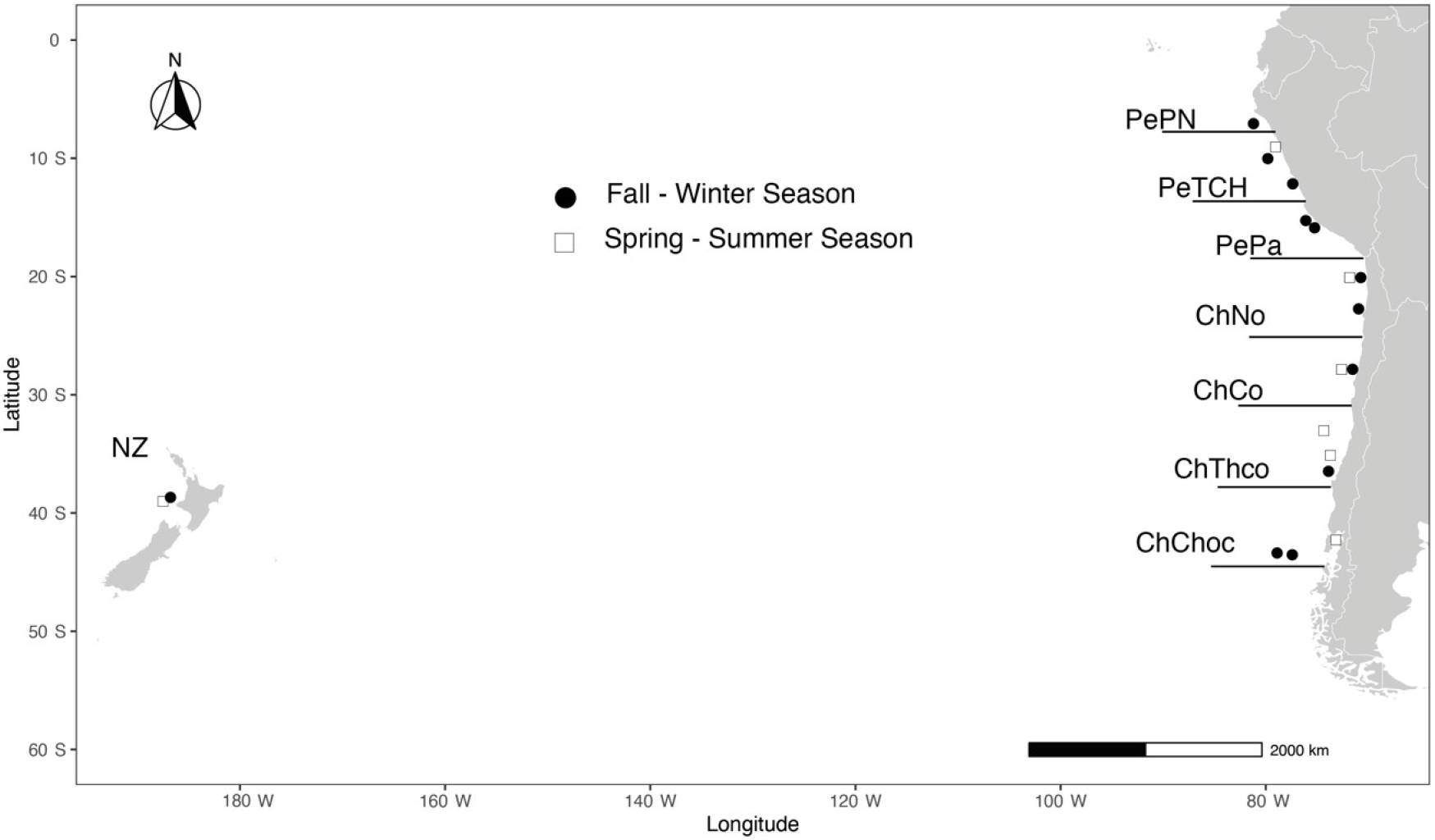
Sampling locations within areas in Fall-Winter and Spring-Summer for *Trachurus murphyi*

### 2.2. Laboratory procedures

For all sampled individuals, we obtained a piece of muscular tissue which was preserved in 96% ethanol for subsequent analyses. The extraction and isolation of the total genomic DNA was obtained using a salting-out protocol as described by Jowett (1986). We assessed the quality and quantity of each DNA purification with an Eppendorf biophotometer®, diluting the template DNA to 20 ng/µL for further PCR amplification. We amplified ten microsatellite loci where three correspond to heterologous loci from *Trachurus trachurus* (Tt29, Tt62, and Tt133; Kasapidis and Magoulas, 2008), and seven correspond to specie-specific loci from *T. murphyi* (Tmur A101, Tmur A104, Tmur A115, Tmur B2, Tmur B6, TmurB104, and Tmur C4; Canales-Aguirre et al., 2010a). The concentrations of reagents for PCR for the heterologous loci were as follows: 1.5 mM of MgCl_2_, 0.2 mM of dNTP’s, 0.2 µM of each primer (3’-end of the forward primer included con dye), and 0.1 U/µL of Taq ADN polymerase (Invitrogen®). The thermocycler conditions were similar to those described in Kasapidis & Magoulas (2008), except that we standardized to 58º C the annealing temperature for all loci. For the species-specific loci, the concentrations of reagents for PCR and thermocycler conditions were conducted following Canales-Aguirre et al., (2010a). All PCRs were conducted using a PTC-200 (MJ-Research®) thermocycler, and we included both positive and negative controls. Amplicons were visualized with an ethidium-bromide-stained gel. A fragment analysis was conducted for PCR products on an ABI 3330 DNA sequencer, and we scored alleles using Peak ScannerTM software v1.0 (www.appliedbiosystems.com/peakscanner), with GS500 as the internal size standard.

### 2.3. Analyses procedures

For summary statistics of genetic diversity, we calculated the total number of alleles (N_A_), the expected (H_E_), and observed (H_O_) heterozygosity for each locus and area using GENALEX v6.5 software (Peakall and Smouse, 2012). We also assessed deviations from Hardy-Weinberg equilibrium and linkage-disequilibrium using ARLEQUIN v3.1 (Excoffier and Lischer, 2010) and GENEPOP 3.1 (Raymond and Rousset, 1995; Rousset, 2008), respectively. We estimated pairwise FST comparisons and a hierarchically AMOVA using ARLEQUIN to test spatial genetic differentiation. The pairwise F_ST_ was conducted between areas within the season, and the significant statistical differences were derived from 10,100 permutations. We applied a sequential Bonferroni correction for multiple comparisons as needed (Rice, 1989), where the adjusted p-value was obtained from α’= 0.05 / (pop x loci). Additionally, an analysis of molecular variance (AMOVA) was performed to test different hypotheses (spatial groupings) of population differentiation that have been obtained through non-genetic approaches. The hypotheses tested were: a) Panmixia (genetic homogeneity between sites); b) Two separate populations in the Southeastern Pacific Ocean as suggested by non-genetics approaches (e.g., George-Nascimento, 2000; Hernández et al., 1998; Serra, 1991); c) Three separate populations considering two in the Southeastern Pacific Ocean and one in Southwestern Pacific Ocean (i.e., New Zealand). We also conducted an AMOVA to estimate if there is a temporal structure among seasons. Finally, we used all previously published genetic information to compare the expected heterozygosity estimated in this study and previous research, considering the sampling collection year and loci tested. We used results from Cárdenas et al., (2009) and Ferrada Fuentes et al., (2023), which used similar geographical areas and loci. The expected heterozygosity was used because it is less sensitive to sample size imbalance than observed heterozygosity (Ritland, 1996).

## 3. Results

### 3.1. Analyses procedures

The summary statistics varied depending on season and area (Table 1). For the fall-winter season, N_A_ ranged from 8.2 to 20.,1 while for the spring-summer season, from 11.3 to 16.9 alleles by locus. Overall, N_A_ in spring-summer was larger than in the fall-winter season. The observed heterozygosity for the fall-winter season ranged from 0.572 to 0.764, while for the spring-summer season from 0.529 to 0.74. The expected heterozygosity for the fall-winter season ranged from 0.701 to 0.839, while for the spring-summer season from 0.633 to 0.832.

**Table 1.**
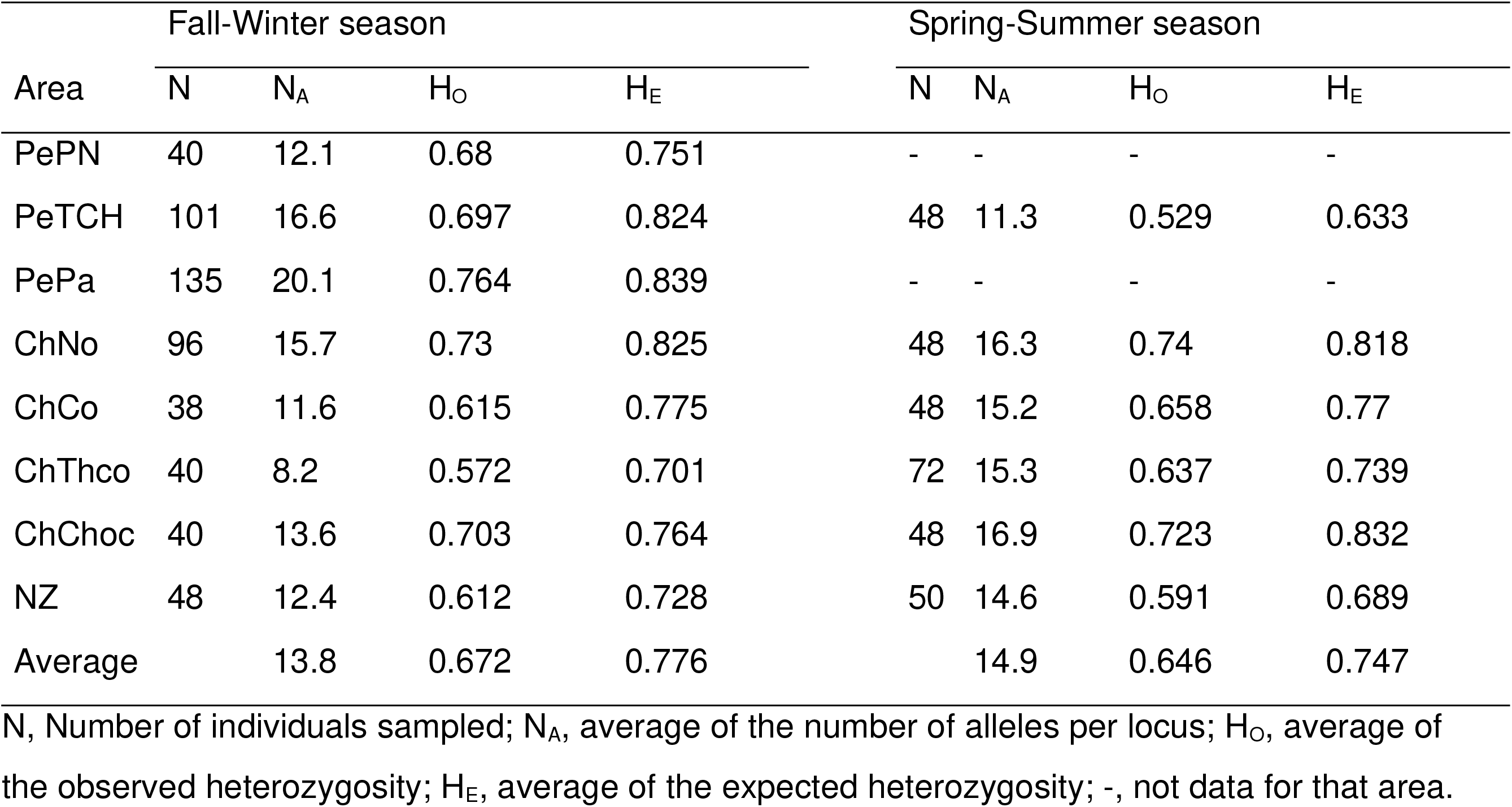
Mean summary statistics for genetic diversity by area in fall-winter and spring-summer inferred for *Trachurus murphyi*.

Pairwise F_ST_ showed a consistent negative and non-significative (p>0.05) F_ST_ pattern, except in the spring-summer season where ChNo-NZ and ChThco-NZ showed a p-value < 0.05 (Table 2). Negative values estimated for fall-winter and spring-summer seasons correspond to F_ST_ equal to zero, indicating that the variation is higher within than among populations.

**Table 2.** Pairwise F_ST_ indices estimated by areas in fall-winter and spring-summer inferred for *Trachurus murphyi*.

|  | Fall-Winter season |  |  |  |  |  |  |  | Spring-Summer season |  |  |  |  |  |  |  |
| --- | --- | --- | --- | --- | --- | --- | --- | --- | --- | --- | --- | --- | --- | --- | --- | --- |
|  | PePN | PeTCH | PePa | ChNo | ChCo | ChThco | ChChoc | NZ | PePN | PeTCH | PePa | ChNo | ChCo | ChThco | ChChoc | NZ |
| PePN | * | 1 | 1 | 1 | 0.997 | 1 | 1 | 1 | * | - | - | - | - | - | - | - |
| PeTCH | -0.086 | * | 1 | 1 | 1 | 1 | 1 | 1 | - | * | - | 1 | 1 | 1 | 1 | 0.542 |
| PePa | -0.054 | -0.014 | * | 1 | 1 | 1 | 1 | 1 | - | - | * | - | - | - | - | - |
| ChNo | -0.129 | -0.020 | -0.018 | * | 1 | 1 | 1 | 1 | - | -0.037 | - | * | 0.412 | 0.221 | 1 | **0.009** |
| ChCo | -0.033 | -0.206 | -0.147 | -0.244 | * | 1 | 1 | 1 | - | -0.046 | - | -0.001 | * | 0.170 | 0.997 | 0.997 |
| ChThco | -0.236 | -0.463 | -0.337 | -0.459 | -0.161 | * | 1 | 1 | - | -0.154 | - | 0.002 | 0.003 | * | 1 | **0** |
| ChChoc | -0.055 | -0.058 | -0.042 | -0.044 | -0.122 | -0.394 | * | 1 | - | -0.124 | - | -0.042 | -0.037 | -0.04 | * | 1 |
| NZ | -0.187 | -0.087 | -0.069 | -0.076 | -0.341 | -0.705 | -0.162 | * | - | 0 | - | 0.010 | -0.023 | 0.022 | -0.037 | * |

The bold values correspond to significant $p$ -values $< 0.05$

The hierarchical analysis of molecular variance for several hypotheses did not show significant differences within each season (Table 3). Similarly to pairwise comparisons, the F_ST_, F_SC_, and F_CT_ in the AMOVA were negative. Testing non-differentiation between the fall-winter and the spring-summer seasons in the AMOVA, we obtained an F_ST_ = -0.01855 (p-value= 1).

**Table 3.** Hierarchical analysis of molecular variance (AMOVA) to test for spatial variation by areas in fall-winter and spring-summer inferred for *Trachurus murphyi*.

| Nº Groups | Aggrupation | Fall-Winter season |  |  |  |  |  | Spring-Summer season |  |  |  |  |  |
| --- | --- | --- | --- | --- | --- | --- | --- | --- | --- | --- | --- | --- | --- |
| | | $F_{ST}$ | p-value | $F_{SC}$ | p-value | $F_{CT}$ | p-value | $F_{ST}$ | p-value | $F_{SC}$ | p-value | $F_{CT}$ | p-value |
| Three | Perú v/s Chile v/s NZ | -0.103 | 1 | -0.103 | 1 | 0 | 0.689 | -0.007 | 1 | -0.061 | 1 | 0.05 | 0.133 |
| Two | Perú+ChNo v/s Chile | -0.14 | 1 | -0.61 | 1 | -0.074 | 0.973 | -0.031 | 1 | -0.026 | 1 | -0.005 | 0.604 |
| One | Panmixis | -0.103 | 1 | - | - | - | - | -0.030 | 1 | - | - | - | - |

Perú includes PePN, PeTCH, and PePa; Chile includes ChNo, ChCo, ChThco, and ChChoc; NZ is New Zealand. Perú+ChNo includes all areas in Perú and only the north of Chile (i.e., ChNo).

The comparisons of the expected heterozygosity among studies showed qualitative differences according to the season in this study, the year of the sampling location, and the locus tested (Table 4). Among studies, loci that did not vary considerably (< 0.1 magnitude) were TmurA104, TmurA115, TmurC4, and Tt62. The loci TmurB6, TmurA101, Tt29, and Tt133 showed a qualitative decline in the H_E_ during the spring-summer season from 2007 to 2012. Conversely, H_E_ qualitatively lows in the fall-winter season were TmurB104 and TmurB2.

**Table 4.** Expected heterozygosity comparisons for this study and previous research, considering the sampling collection year and loci tested.

| Season / Previous studies | Year collection | Expected heterozygosity |  |  |  |  |  |  |  |  |  |
| --- | --- | --- | --- | --- | --- | --- | --- | --- | --- | --- | --- |
|  |  | TmurB6 | TmurA104 | TmurA115 | TmurC4 | TmurA101 | TmurB104 | TmurB2 | Tt29 | Tt62 | Tt133 |
| Ferrada Fuentes et al. (2023) | 2007 | - | 0.943 | 0.95 | - | 0.927 | - | - | 0.747 | 0.855 | 0.861 |
| Cárdenas et al. (2009) | 2009 | - | - | - | - | - | - | - | 0.76 | 0.83 | 0.87 |
| Fall-Winter | 2011 | 0.782 | 0.93 | 0.916 | 0.82 | 0.871 | 0.521 | 0.552 | 0.758 | 0.851 | 0.758 |
| Spring-Summer | 2011-2012 | 0.444 | 0.942 | 0.945 | 0.819 | 0.779 | 0.648 | 0.704 | 0.634 | 0.865 | 0.687 |

## 4. Discussion

Research on *Trachurus murphyi* populations in the South Pacific Ocean has mostly concentrated on fisheries data from the non-reproductive season. Previous studies have revealed genetic homogeneity across this region, which aligns with our findings. Interestingly, we also noticed a consistent pattern in the fall-winter and the spring-summer seasons. This could indicate a regular movement of individuals between feeding and spawning areas along the coast of the Southeastern Pacific Ocean, potentially leading to genetic structuring. More research is needed, especially regarding microsatellite loci, to fully understand this phenomenon. From 2007 to 2012, a noteworthy aspect was the identification of four loci with diminished levels of expected heterozygosity during the spring-summer season and two during the fall-winter seasons. These findings suggest that these specific genetic markers may serve as effective indicators for monitoring changes in genetic diversity. This research calls into question the idea that differences in time, such as spawning and feeding seasons, may result in distinct temporal population structures.

Our study has shown that there is a lack of spatial population structure both within each season (Fall-Winter and Spring-Summer) and between them, with low pairwise F_ST_ values and AMOVA results (p < 0.05). This supports previous research that found marine fish populations often have F_ST_ values below 0.01 (Ward et al., 1994). These results are also consistent with studies conducted on *T. murphyi* in the same region, using different markers such as isozymes (Galleguillos and Torres, 1998), RFLP-PCR (Sepúlveda et al., 1996), mitochondrial DNA, and heterologous and species-specific microsatellites (Canales-Aguirre et al., 2010a; Cárdenas et al., 2009; Ferrada Fuentes et al., 2023; Galleguillos et al., 2012). In the Southeastern Pacific Ocean, these findings are not uncommon and have been previously observed in other species within the same region. Additionally, a similar trend of genetic similarity has been noticed in other marine species distributed across this area, including *Engraulis ringens, Strangomera bentinki, Sprattus fuegensis, Genypterus blacodes*, and *Dissostichus eleginoides* (Canales-Aguirre et al., 2018, 2016, 2010b; Ferrada et al., 2002; Galleguillos et al., 1997). These findings suggest that geographical barriers may not significantly drive divergence within this particular region.

Marine fish species, such as *T. murphyi*, typically possess several characteristics that contribute to maintaining genetic similarity among populations. These include large numbers of individuals, a tendency to reproduce and develop in the open ocean, high reproductive rates, and the ability to migrate. These factors have been extensively studied and have been found to play a crucial role in minimizing genetic divergence among populations (Serra, 1991; Wirgin and Waldman, 2005). The population dynamics of *T. murphyi* are heavily influenced by factors such as population size and the formation of reproductive aggregations, which occur in the Southeastern Pacific Ocean, where the species is ubiquitously found. These aggregations are known to occur throughout the species’ entire range (Gorbunova et al., 1985), indicating that the species is highly abundant and displays extensive migratory behaviors. Additionally, current research has revealed that *T. murphyi* prefers to inhabit areas with specific temperature ranges (15 -18 oC Cubillos et al., 2008). Different intensities of El Niño events strongly impact the habitat patterns of commercially valuable species off the Chilean coast, altering suitable areas and distribution (Feng et al., 2022). This could potentially lead to genetic mixing across the entire geographic range of the species. Our research shows no genetic differentiation in *T. murphyi* over space and time throughout its distribution. Despite considerable distances between sample locations, the ability of a small number of individuals to migrate and successfully reproduce is enough to maintain genetic homogeneity within the population.

Upon analysis, the ten loci examined displayed polymorphism throughout their geographical range. When comparing the mean summary statistics of genetic diversity to those reported by DeWoody & Avise (2000) for marine fish, we noted that the number of alleles for fall-winter (N_A_= 13.8) and spring-summer (N_A_= 14.9) samples fell below the average (N_A_=20.4). Similarly, the observed averages for heterozygosity in fall-winter (0.672) and spring-summer (0.646) were lower than the averages reported for other marine species (H_O_= 0.790), indicating lower levels of genetic diversity in our study. The results of our study on *T. murphyi* have shown varying outcomes compared to previous research. While our estimation of expected heterozygosity (H_E_) using the same heterologous loci was generally low in the spring-summer season (2 out of 3 loci) and showed no distinct pattern in the fall-winter season (1 out of 3 loci), it is important to acknowledge the limited number of shared heterologous loci available for comparison (Cárdenas et al., 2009). Furthermore, upon examining the findings of Ferrada Fuentes et al. (2023), we discovered that over 60% of the loci displayed low H_E_ in both seasons, with 5 out of 6 comparisons exhibiting lower values in the spring-summer season and 4 out of 6 in the fall-winter season. This highlights the need for caution in drawing definitive conclusions, given the scarcity of shared heterologous or specific loci for accurate comparison.

Our research uncovered fluctuations in expected heterozygosity at specific loci over time. Notably, expected heterozygosity, determined by allele frequencies, is more reliable than observed heterozygosity regarding sample size bias. This is because observed heterozygosity relies on the genotypes of individual samples (Ritland, 1996). Previous studies on the population genetics of *T. murphyi* have mostly relied on data collected during a single season. This has left gaps in our understanding of temporal population trends, especially during reproductive and non-reproductive phases. The motivation for this method stems from the idea that migration to certain reproductive areas could result in greater variety and changes in genetic diversity between seasons. Waples (1998) proposes that sampling during the reproductive season is ideal for understanding population genetic patterns and reducing bias from related individuals (Allendorf and Phelps, 1981; Waples, 1998). While a meta-analysis of previous databases was not conducted, examining the expected heterozygosity over time offered valuable insight. Focusing on the fall-winter and spring-summer seasons provided a unique outlook on temporal diversity within a given year. Although variations in expected heterozygosity in certain loci from 2007 to 2012 indicated their potential for monitoring genetic diversity, overall differences were not significant enough to suggest a temporal population structure. However, some loci, such as TmurB6, did exhibit qualitative variation across both seasons.

Although this study’s results are consistent with prior research, it is crucial to consider certain shortcomings. Using samples solely from active fishery vessels introduces potential variations in sampling locations, resulting in the need for wider geographical ranges rather than precise sites. As a result, certain regions in Perú were inaccessible for sampling, which may affect comparisons.

Moreover, previous studies on *T. murphyi* have utilized partially overlapping loci in their microsatellite analyses, posing challenges for direct comparisons of expected heterozygosity. Despite these limitations, this study has incorporated data from a minimum of three distinct years, covering from 2007 to 2012, to support the study’s temporal reliability. The F_ST_ values are concerning due to the unequal sample sizes, highlighting potential issues with sample size. Our data sets ranged from 38 to 135 for the fall-winter samples and 48 to 72 for the spring-summer samples. Despite these negative F_ST_ values, reporting them as F_ST_ = 0 is customary, indicating a greater genetic variation within populations rather than between them (Weir and Cockerham, 1984). However, using ARLEQUIN software, which incorporates Weir and Cockerham’s methodology, addresses this issue, but the negative values still point to a possible inadequacy in sample size (Gerlach et al., 2010). Therefore, further sampling efforts may be necessary to ensure balanced sample sizes.

To gain a comprehensive understanding of genetic differences, future analyses must prioritize the utilization of single nucleotide polymorphisms (SNPs) obtained through reduced-representation sequencing techniques (Campbell et al., 2018), such as RADseq, ddRAD, or DARTseq (Kilian et al., 2012; Miller et al., 2007; Peterson et al., 2012). These methods have demonstrated their effectiveness in distinguishing genetic variations, particularly in species with relatively shallow genetic structures and high rates of gene flow, even on a small geographical scale (Canales-Aguirre et al., 2022). These advanced techniques allow researchers to explore hypotheses surrounding neutral and adaptive genetic variation. For instance, studying neutral loci may uncover patterns of isolation by distance (e.g., Drinan et al., 2018), a particularly relevant consideration given the expansive range of *T. murphyi* throughout the Southeastern Pacific Ocean and Southwestern Pacific Ocean (∼10,000 km). Adaptive loci could offer valuable insights into genetic differentiation linked to local adaptation in the species’ most extreme distribution areas. In addition, analyzing the entire genome over multiple years of data collection could further reinforce and validate the conclusions drawn from this study.

Our research unequivocally backs the concept that *T. murphyi* displays a stable, homogenous population genetic trend in all areas it inhabits, with no difference seen over time. By investigating specimens gathered during mating and non-mating seasons over multiple years in identical settings, we have illuminated a crucial aspect of temporal distinction, closing a significant void in current literature. Finally, the loci with reduced heterozygosity that have been identified could be utilized to detect changes in genetic variability in this species under climate change or fishery pressure scenarios.

## Acknowledgments

We extend our gratitude to the scientific observers from Instituto de Fomento Pesquero (IFOP) in Chile, Instituto del Mar del Perú (IMARPE) in Perú, and the Ministry for Primary Industries in New Zealand, facilitated by the National Institute of Water and Atmospheric Research (NIWA), for their invaluable assistance in collecting the Chilean jack mackerel samples. Additionally, we thank Mariam Dib and Andrea Barrera for their invaluable contributions to the laboratory procedures. Cristian B Canales-Aguirre receives funding from ANID – Millennium Science Initiative Program – NCN2021-056.

## Funding

This research was funded by Fondo de Investigación Pesquera y Acuicultura, grant number FIPA 2010-18, from the Chilean Government.

## CRediT authorship contribution statement

**Cristian B. Canales-Aguirre**: Conceptualization, Data curation, Formal analysis, Investigation, Writing – original draft, Writing – review & editing; **Sandra Ferrada Fuentes**: Conceptualization, Data curation, Formal analysis, Investigation, Visualization, Writing – review & editing; **Ricardo Galleguillos**: Conceptualization, Funding acquisition, Investigation, Project administration, Resources, Writing – review & editing.

## Declaration of Competing Interest

The authors declare that they have no known competing financial interests or personal relationships that could have appeared to influence the work reported in this paper.

## Data availability

The data presented in this study are available at https://github.com/Canales-AguirreCB/Canales-Aguirre_etal_2024_FishRes.

## Notes

### Competing Interest Statement

The authors have declared no competing interest.

